# Models for Evaluation of Biological Age Based on Multi-omics

**DOI:** 10.1101/2021.12.13.472382

**Authors:** Ce Huang, Yingqian Zhu, Fengjiao Huo, Shengyu Feng, Xue Gong, Hua Jiang, Zhongmin Liu, Hailiang Liu

## Abstract

Aging is a complex process of systemic degeneration at multiple cellular and tissue levels that has a complex mechanism. Individuals age at different rates, and there is a high degree of aging heterogeneity. Therefore, it is inaccurate to judge an individual’s degree of aging by their chronological age. We performed transcriptome-focused multi-omics analyses (including transcriptomics, DNA methylation, proteomics, cytokine and telomere analysis, and single-cell transcriptome sequencing) on 139 healthy individuals aged 23 to 88 years. We systematically analyzed linear and non-linear changes in gene expression throughout aging. Genes with similar expression trajectories were enriched in similar biological pathways, including the PI3K-AKT signaling pathway and the inflammatory response. Through DNA methylation detection, we found that the expression of the top genes correlated with age was affected by methylation in the gene promoter region. These genes had no significant correlation with the expression of downstream proteins, but they were enriched in PI3K-AKT-associated proteins. Single-cell transcriptome sequencing showed that the expression of these genes did not significantly change in different cell subtypes, which proves that the gene expression changes were caused by internal age-related cellular changes rather than cell composition changes. We designed a transcriptome age clock and a methylation age clock using a set of 787 genes. Our models can accurately predict age with a mean absolute error (MAE) of 5.203 and 3.28, respectively, which is better than previously established aging models. The accuracy of our model was further verified by the detection of telomeres, which can identify accelerated aging of individuals.

---

Aging is a biological process that gradually leads to a decline in physiological functions and ultimately leads to organ failure followed by death. In this process, individuals tend to age at different rates and in different ways. It is difficult to measure the degree of aging based on the time elapsed since birth, known as an individual’s chronological age, as this fails to represent the functional status of different individuals. In contrast, biological age quantified by biomarkers can provide an accurate indication of the aging process. Therefore, biomarkers that can predict the age and life expectancy of individuals is an important prerequisite for studying the mechanisms of aging.

At present, an increasing number of researchers use omics methods in the fields of aging, including transcriptomics, metabolomics, proteomics, and epigenetics. Epigenetic modifications or dynamic changes to the expression products of the genome with advancing age are found in multiple omics models. These modifications and expression products can constitute an accurate dataset for predicting the age of an individual. At present, the most common predictor is based on DNA methylation data, proposed by Horvath and known as “epigenetic clocks” (Hannum *et al*. 2013; Horvath 2013; Levine *et al*. 2018), which can estimate age based on DNA methylation (DNAm) in specific tissues or predict the age of an individual independently of tissues. The transcriptome is regulated by DNA methylation. At the same time, the transcriptome itself is an important regulatory component in the cell life cycle. The relationship between transcriptome levels and aging has recently been of great interest to researchers. Based on microarray chip data of whole blood samples, an aging clock of R^2^ = 0.6 and mean absolute error (MAE) = 6.24 years was established (Peters *et al*. 2015), but the model performed poorly (Fleischer *et al*. 2018). This was related to the limitations of microarray data, including background noise and a limited detection range. Recently, with the development of second-generation transcriptome sequencing technology, we can obtain more accurate and comprehensive transcript expression data through RNA-seq. Transcriptome expression data from RNA-seq on human dermal fibroblasts cultured *in vitro* were used to develop another aging prediction model (R^2^ = 0.81, MAE = 7.7). However, because of the difficulties and uncertainties in obtaining and culturing human dermal fibroblasts, this method is not suitable as a routine index for clinical diagnosis. Blood samples are mainly collected through venipuncture, which is much less invasive, has a wide range of applications, is convenient, and is the best source for research on aging markers. The DNA methylation clock has been established mainly by whole blood and peripheral blood mononuclear cells. At the same time, studies based on plasma proteins in the blood have found that 1,379 plasma proteins fluctuate with age, and thus a plasma protein aging model (Pearson = 0.97) has been established that performs better than the epigenetic clock model (Lehallier *et al*. 2019). However, compared with RNA-seq, proteomics detection still has technical complexity and uncertainty. It has been proven that RNA-seq data can accurately predict the transcriptional aging clock of individual age. However, there have been no relevant RNA-seq studies on human blood samples to date. Thus, we performed this study, aiming to establish a model of biological age that can accurately predict the functional status based on the gene expression of healthy individuals.

## Results

### Differential genes appear to increase with age

Aging causes the function of the blood circulatory system to decline and destroys the biological processes that maintain health. At present, we still lack an understanding of the differences in the incidence and rate of aging of the blood system, especially the peripheral blood mononuclear cells (PBMC). Therefore, we selected a population of 139 relatively healthy individuals (ages 23-88 years old; median, 53 years old) by multiple clinical tests (including 12 chronic diseases) (Figure 1A, Extended data Table S1). We performed a paired differential gene expression analysis to determine when differentially expressed genes (DEGs) appeared and whether they persisted with age, 113 samples were transcriptome sequenced. First, we screened 21,024 genes, removed genes with low expression levels (read count > 30), and got a total of 13,571 genes. After that, we grouped all samples into seven groups based on age, the 20-year-old group (20–29 years old), the 30-year-old group (30–39 years old), the 40-year-old group (40–49 years old), the 50-year-old group (50–59 years old), the 60-year-old group (60–69 years old), the 70-year-old group (70–79 years old), and the 80-year-old group (over 80 years old). All pairwise comparisons took the 20-year-old group as the control. The results show that compared with 30- and 40-year-olds, the number of DEGs were significantly increased in the 50-80-year-old groups (Figure 1B), which indicates the progress of gene expression such that a significant change in genes will only occur after sufficient time. In the 70-year-old age group, it seems that the effect of aging on gene expression changes has declined. This may be partly due to the relatively large variance caused by the large age distribution of this group. With increasing age, pathways such as focal adhesion, ECM-receptor interaction, and the PI3k-Akt signaling pathway was enriched (Figure 1B). As shown in Figure 1C, the numbers of upregulated genes in each age group were significantly less than the numbers of downregulated genes. In addition, we found not only that the number of differential genes was small at the ages of 30 and 40, but also these differential genes did not persist. Only *SERPINF1* remained downregulated through the entire age group (Figure 1C). From the age of 50, a relatively conservative set of core aging characteristics gradually became maintained (Figure S1A). For detailed genetic information, see Extended data Table S2. Among them, 43 genes continued to be upregulated (Figure 1C). Genes that continued to be upregulated with age were mainly related to inflammation, such as the top differential genes inflammasome (NLR family CARD domain containing 4, *NLRC4*) and sialic acid-binding immunoglobulin-like lectin 15 (*SIGLEC15*). NLRC4 forms cytosolic multiprotein complexes that sense microbial infections or host cell damage, triggering cytokine production and a proinflammatory form of cell death called pyroptosis (Andrade and Zamboni 2020). NLRC4 can directly bind to caspase-1 through its CARD-CARD domain, resulting in caspase-1 activation with no need for autoproteolysis (Poyet *et al*. 2001; Broz *et al*. 2010). SIGLEC15 is overexpressed in various cancers and is a novel antitumor target comparable to programmed cell death 1 ligand 1 (PD-L1) (Wang *et al*. 2019; Li *et al*. 2020). After the age of 50, 294 genes continued to be downregulated (Figure 1C & Figure S1B), among which phosphodiesterase 4D (PDE4D) is a newly discovered candidate target for Alzheimer’s disease, a neurodegenerative disease closely related to age. PDE4D has been found to be related to the cognitive function of the dorsolateral prefrontal cortex (dlPFC) of primates. As the dlPFC ages, the expression of PDE4D will decrease. Age-related loss of PDE4D may thus contribute to the specific vulnerability of the frontal cortex to degeneration in Alzheimer’s disease and may play a critical role in normal cAMP regulation (Datta *et al*. 2020; Leslie *et al*. 2020) (Figure 1D).

**Figure 1.**
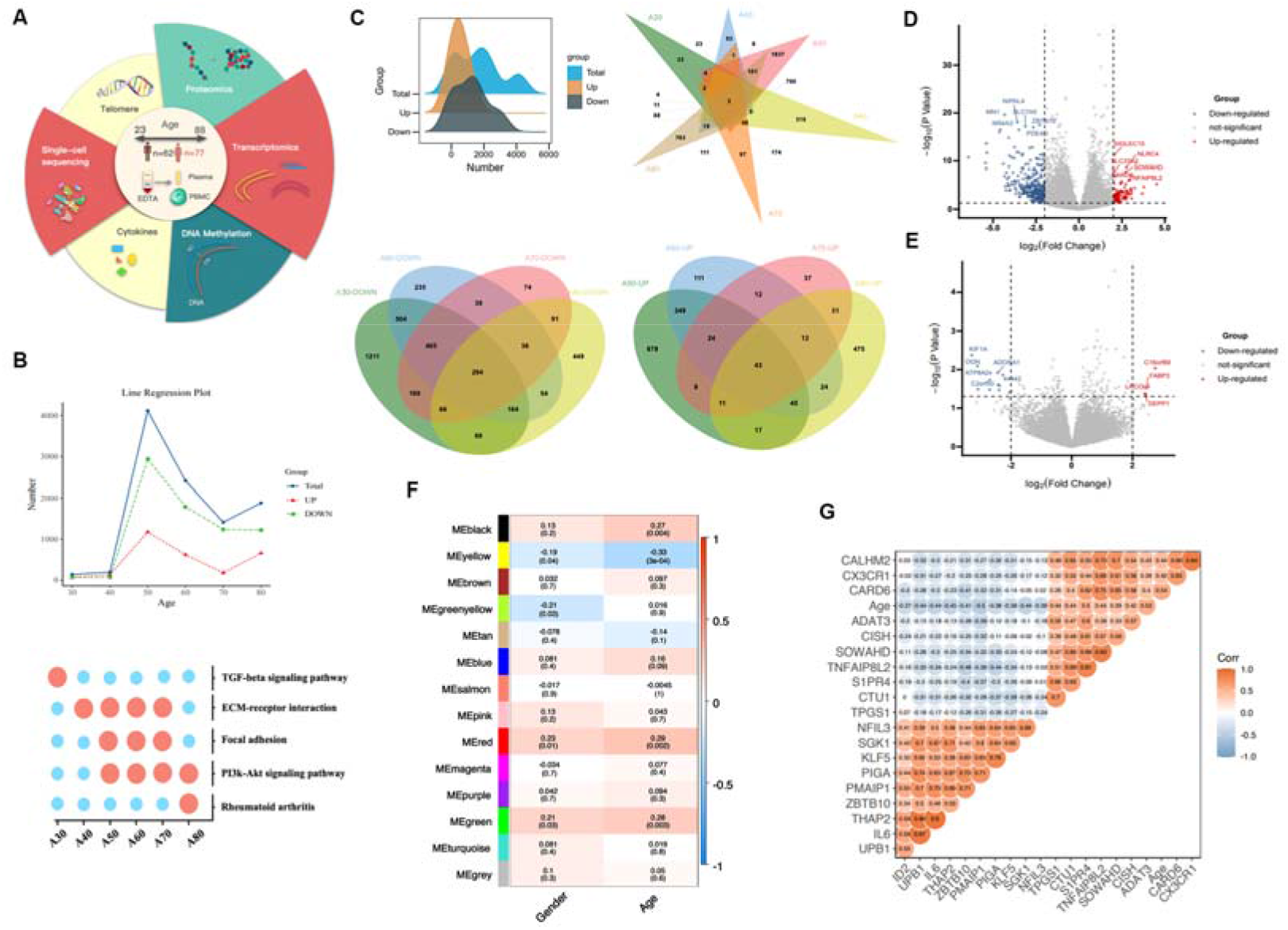
Pairwise differential expression across ages. **A** Schematic of the experiment. Blood samples were collected from males (n = 62, 24–88 years old) and females (n = 77, 23–88 years old), the PBMC and plasma were separated, and six omics tests were performed. **B** With the 20-year-old group as the control, the number of differential genes for each age group compared in pairs are shown: blue, red, and green represent the total differential genes, the upregulated differential genes, and the downregulated differential genes, respectively. The DEGs enrichment pathways (database source: KEGG). **C** Ridge map showing the density distribution of the differential genes, and three Venn diagrams showing the distribution of the total differential genes in the six age groups (30, 40, 50, 60, 70, and 80 years old). The difference is upregulated. The distribution of genes in four age groups and the distribution of the downregulated differential genes in four age groups (50, 60, 70, and 80 years old). **D** Volcano map of conservatively different genes between ages (based on the logFC and p-values of the 20-year-old vs. 80-year-old age groups, logFC > 2, p < 0.05). **E** Volcano map of conservatively different genes between sexes (based on the logFC and p-values of the 80-year-old group, logFC > 2, p < 0.05). **F** Weighted gene coexpression network analysis (WGCNA) and age and sex correlation analysis, showing, respectively, the correlation coefficient (Pearson) and correlation p-value. **G** Differential genes shared between the four age groups (50, 60, 70, and 80 years old) shown in a correlation diagram of the top 20 differential genes.

Sex is also an important factor in the aging process. In proteomic studies, it was found that aging across cohorts and sexes in the proteome maintains a high degree of consistency. In our study, the analysis of differential gene expression between sexes in each age group showed that the number of differential genes caused by sex is also numerous (Figure S1C). However, the DEGs are generally very different (Figure 1D and Figure 1E), only in the 60-year-old group was there a large overlap between the two. At the same time, according to the weighted gene coexpression network analysis (WGCNA) results (prior to carrying out trend analysis, COMBAT was used to adjust for differences between RNA-Seq Batch 1, Batch 2 and Batch 3) (Figure 1F & Figure S1D), the age-related modules (such as the MEred and MEyellow modules) are highly correlated with sex. Similar results have been seen in other studies, genomic differences between the sexes increase after age 65, with men having higher innate and pro-inflammatory activity and lower adaptive activity (Márquez *et al*. 2020). Two-thirds of the proteins that changed with age were observed to also change with sex (895 of 1,379 proteins) in plasma proteomics (Lehallier *et al*. 2019). Therefore, sex differences will affect our observations herein on the transcriptome of the aging process. Finally, through correlation analysis (Figure 1G), the genes that were conservatively upregulated in aging were positively correlated from ages 23 to 88, while the downregulated genes were negatively correlated with age.

### Gene expression changes with age

Paired DEG comparisons are inherently limited, and our data can be used to check and observe gene expression dynamics throughout the life cycle. We first searched for genes with common expression trajectories throughout the life cycle. We visualized 13,571 genes as z-score changes throughout the life cycle, and their overall performance showed obvious fluctuations (Figure 2A). To reduce the complexity of the data, we used unsupervised hierarchical clustering to group genes with similar trajectories (Figure 2B), and so identified eight trajectory clusters. These clusters changed with age and their size ranged from 860 to 2,296 genes (Figure 2C). In addition to the linear mode (cluster 1 and 7), some non-linear trajectories could also be seen (Figure 2C). By correlation (Spearman’s) analysis of gene expression with age in individual patients, 3,956 genes out of 13,571 were significantly correlated with age (p < 0.05), of which 2,667 were positively correlated and 1,289 were negatively correlated (Extended data Table S2). In cluster 1, 963 genes were significantly associated with age, all of which were positive; in cluster 7, 695 genes were significantly associated with age, all of which were negative. Eight clusters were examined for patterns of functional enrichment. In the Kyoto Encyclopedia of Gene and Genomes (KEGG) pathways, seven clusters were enriched for specific biological pathways (q < 0.05; Figure 2C), which indicates that the biological processes are precisely regulated during the life process. For example, PI3K-AKT continues to decline with age (cluster 7), and the cluster also contains anti-inflammatory factors such as IL4 (Figure 2D, Extended data Table S3). In addition, clusters 3, 4, and 5 show similar trajectories, which we termed “N-type fluctuating trajectories”. The top three enriched pathways were all enriched for Herpes simplex virus 1 infection. No related infections were observed on clinical examination of the patients. In general, most genes changed in a non-linear manner throughout the life cycle.

**Figure 2.**
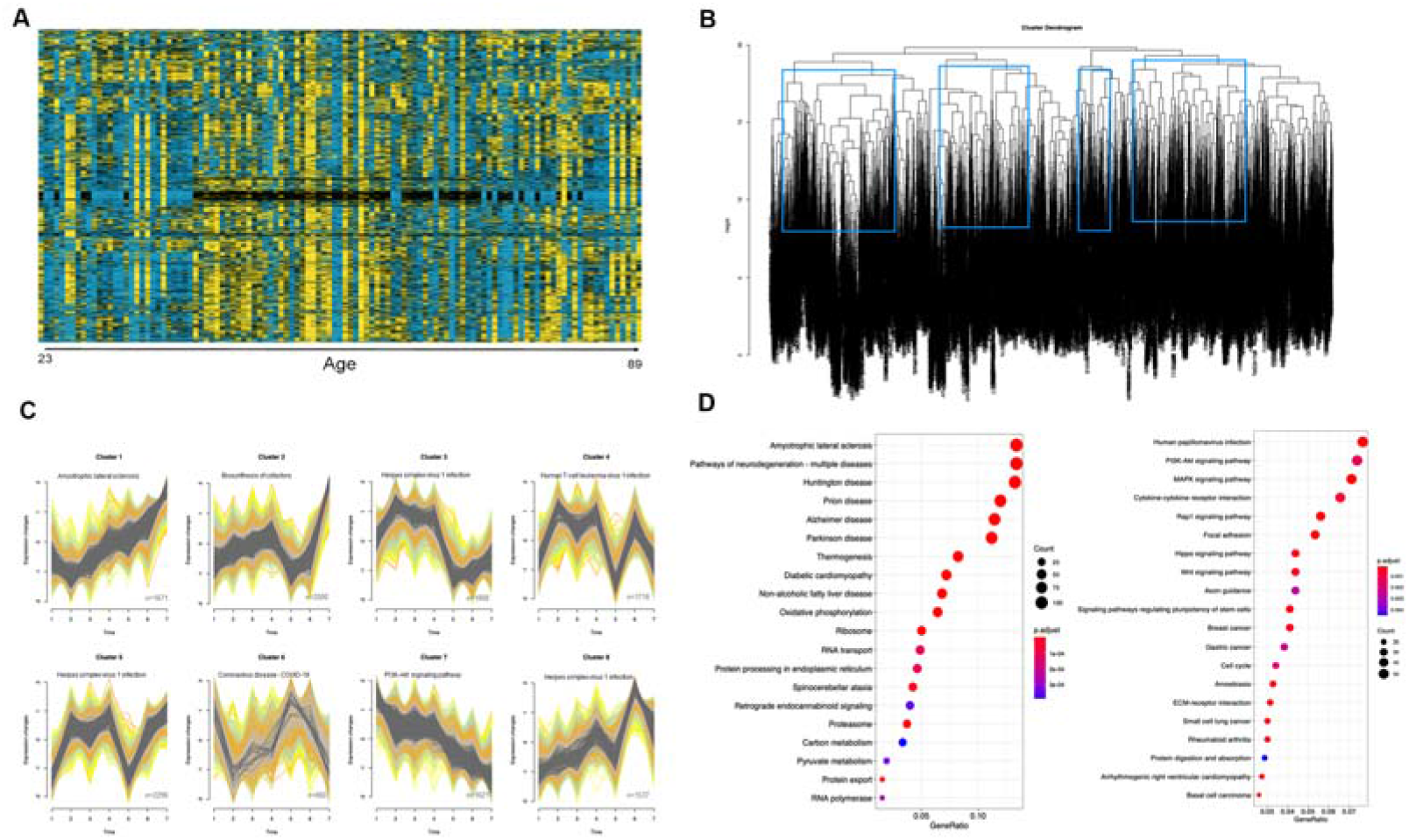
Dynamic analysis of aging gene expression at all ages. **A** Heatmap of gene fluctuation trajectories. Unsupervised hierarchical clustering was used to group genes with similar trajectories. **B** Hierarchical clustering dendrogram. The eight identified clusters are represented by blue boxes. **C** The gene trajectories of the eight identified clusters. The thick line represents the average trajectory of each cluster, and the number of genes in each cluster and the most important enrichment pathway are also marked. The Kyoto Encyclopedia of Genes and Genomes (KEGG) database was used for enrichment. **D** The enrichment pathways of cluster 1 and cluster 7, showing a linear relationship (database source: KEGG).

### The correlation between DNA methylation and gene expression

The level of genomic DNA methylation is highly correlated with age. “Epigenetic clocks” (Horvath 2013) have been established that can estimate the age based on DNAm, either in specific tissues or independently of tissues. These can predict the age and life expectancy of the individual (Levine *et al*. 2018). However, it is not yet known whether these sites regulate the downstream transcriptome, and whether the changes observed in transcription levels with age are related to changes in upstream DNA methylation. In this study, Infinium Methylation EPIC Bead Chip (850K) methylation sequencing was performed on PBMC of 49 healthy individuals aged 23–88. We divided the age distribution into four groups, the 20–35 age group (young, Y), the 40–55 age group (middle, M), the 60–75 age group (young old, YO), and the 80–88 (aged, A) age group. As we did for the transcriptome data, we first performed a paired differential locus analysis to determine when the differential loci appeared and whether they persisted with age. As shown in Figure 3A and Figure S2, the Y and M groups had fewer different sites (8,805), while there were more than 30,000 different sites in the YO and A groups. These differential sites were mostly concentrated in twelve top sites on the chromosome. In the M, YO, and A groups, there were 750 sites with elevated methylation levels and 529 sites with downregulated levels, which were conserved between the M, YO, and A groups. These sites correlated with the results of the transcription composition pair comparison (Figure 3B). We found 18 genes with changes to both the DNA methylation and transcription levels (Figure 3C). These methylation sites were located both in promoter regions and within the bodies of genes. One of these genes, *FLT1*, encoding the cell surface receptor for VEGFA, VEGFB, and PGF, plays an important role in the regulation of angiogenesis, cell survival, cell migration, macrophage function, chemotaxis, and cancer cell invasion. It can promote the proliferation, survival, and angiogenesis of skin cells in adulthood (Luttun *et al*. 2002; Fischer *et al*. 2008; Stefater *et al*. 2013). Another, *FLRT2*, functions in cell adhesion, cell migration, and axon guidance, and mediates cell adhesion by interacting with ADGRL3 and possibly other latrophilins expressed on the surface of adjacent cells (Seiradake *et al*. 2014). In general, however, the number of genes found with changes to both the DNA methylation and transcription levels was extremely small.

**Figure 3.**
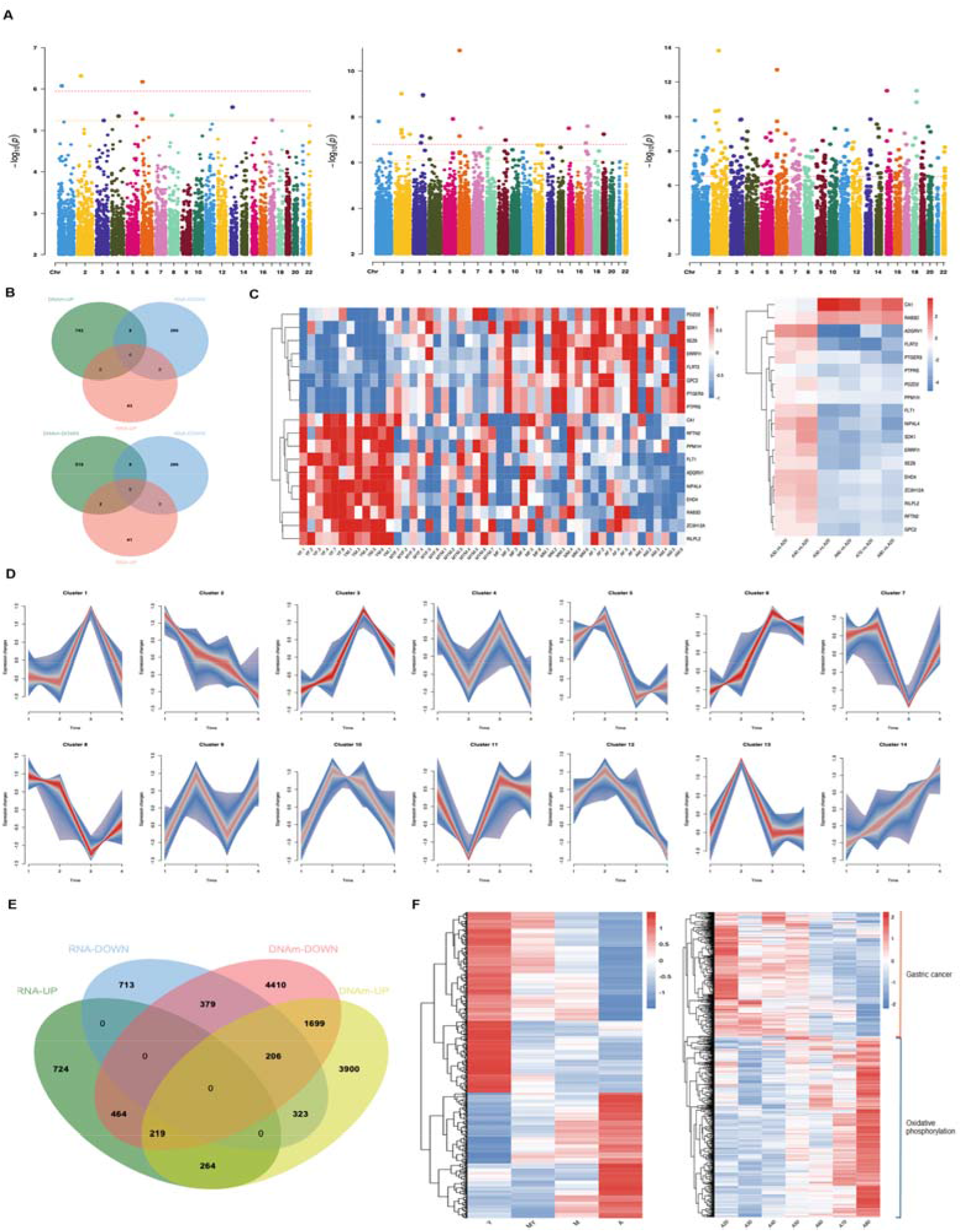
Association analysis of DNA methylation and aging-related gene expression. **A** Manhattan chart showing the information of the differential methylation sites between different ages, with the log_10_ conversion of the difference analysis p-value on the y axis and the chromosome position of each CpG on the x axis. **B** Venn diagram of conservative differential genes (RNA-UP, RNA-DOWN) and conservative differential sites (DNAm-UP, DNAm-DOWN), a total of 18 common genes. **C** Heatmap of the 18 differential genes obtained in Fig B, the left side shows the normalized heatmap of the methylation β values of the 18 differential genes in 49 samples, and the right side shows the 18 differential genes in the paired differential genes. The heatmap on the right shows logFC in the comparisons. **D** The CpG sites trajectory of 14 unbiased clusters. The thick line represents the average trajectory of each cluster, and the number of probes in each cluster is also marked. **E** Venn diagram of the gene expression trajectories of cluster 1 (RNA-UP) and cluster 7 (RNA-DOWN) and the methylation site fluctuation trajectories of cluster 2 (DNAm-DOWN), and cluster 14 (DNAm-UP). **F** Heatmap of the 787 genes obtained in Fig E in the four methylation groups (Y, M, YO, and A, normalized by β value z-scores) and seven transcriptome samples (20, 30, 40, 50, 60, 70, and 80 years old; where FPKM performed z-score normalization).

Next, as for the transcriptome data, we searched for sites with common methylation trajectories throughout the life cycle. We first removed sites located on the sex chromosomes and sites that failed to be annotated to a gene, and then visualized the β values of 150,827 sites located in promoter regions as z-score changes throughout the life cycle, and their overall trajectories (Figure 3D). There were 14 trajectory clusters, which varied with age, ranging in size from 8,164 to 10,740 sites. Cluster 2 and cluster 14 were in a linear pattern. We correlated the genes contained in these sites with clusters 1 and 7 from the transcriptome data, which resulted in 787 genes (Figure 3E, Extended data Table S4). Methylation of the promoter region of genes inhibits gene expression, and so the expression trend of these genes with aging may be caused by DNA methylation in the upstream promoter region. We visualized the β values and expression levels of the 787 genes according to the trajectory, and we found that there was indeed an opposite trend in the DNA methylation and transcription levels. The most enriched pathway of the genes with an upward trend was oxidative phosphorylation (from the KEGG database), and the top pathway of the genes with a downward trend was gastric cancer.

### The correlation between plasma protein and gene expression

A metachronous symbiosis experiment proved that the plasma of young mice can reverse the aging of various tissues and organs of old mice through the blood circulation system (Loffredo *et al*. 2013; Katsimpardi *et al*. 2014; Baht *et al*. 2015; Huang *et al*. 2018). These results indicate that plasma protein is a key regulatory factor in the aging process. Determining the key proteins in the plasma could help us understand the mechanisms of aging. In the proteomics experiment, 24 samples from the cohort were included. Group Y included 12 cases from the 20–30 age group (6 men and 6 women), and Group A included 12 cases from the 80–90 age group (6 men and 6 women). As shown in Figure 4A, with a threshold of fold change (FC) > 1.5 and p < 0.05, a total of 103 differential proteins were obtained, of which 52 were upregulated and 51 were downregulated. As for the transcriptome data, the downregulated proteins were also enriched for the PI3K-AKT signaling pathway (Figure 4B), including IGF1 and CSF1. This pathway also plays an important regulatory role in the entire protein network (Figure 4C). IGF1 is known to be related to longevity, while CSF1 is a cytokine that plays an important role in the regulation of the survival, proliferation, and differentiation of hematopoietic precursor cells, especially monocytes. It promotes the release of proinflammatory chemokines, thereby playing an important role in innate immunity and inflammation. The resulting differential proteins were compared with the linearly related genes, and we found that there was not much overlap. Six genes (*ACTR1A, CST3, LGALS1, LGALS3, OAF*, and *RHOG*) were upregulated at both transcriptomic and proteomic levels, while one gene (*SRC*) was downregulated at both transcriptomic and proteomic levels (Figure 4D).

**Figure 4.**
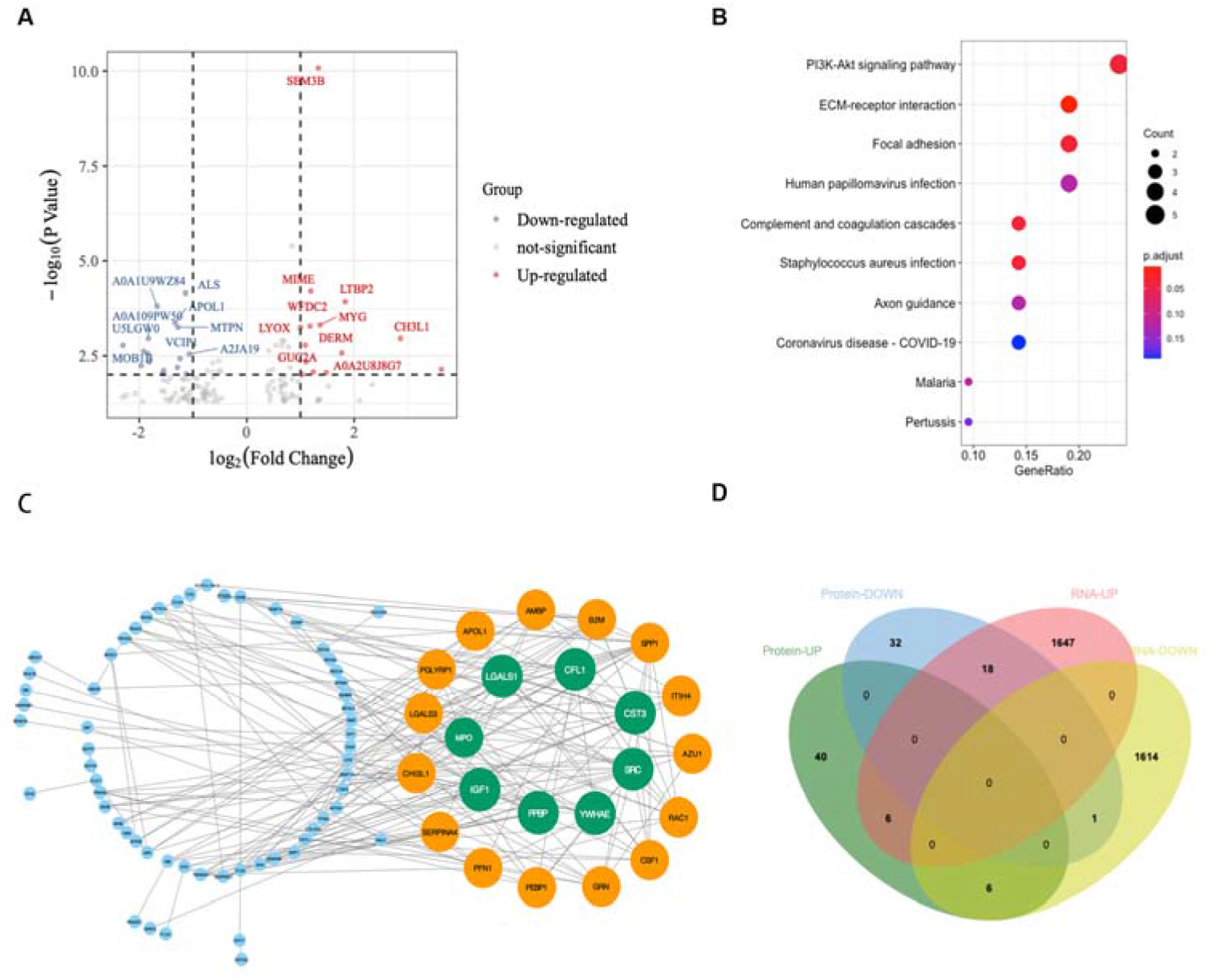
Association analysis of plasma protein and aging-related gene expression. **A** Differential protein volcano diagram between the 20-year-old group (Y group) and the 80-year-old group (A group) (fold change > 1.3, q-value < 0.05). **B** The downregulated differential protein enrichment pathways (database source: KEGG). **C** We performed protein network interaction analysis on 138 differential proteins in STRING, put the output through Cytoscape software, and marked the proteins with the top 23 combined scores in green and yellow. **D** Venn diagram of gene expression trajectory cluster 1 (RNA-UP) and cluster 7 (RNA-DOWN) and the differential proteins (Protein-DOWN, and Protein-UP).

### Single cell sequencing

An important question in bulk RNA transcriptomics is whether the observed changes in gene expression are caused by internal changes in the cells because of age, or by changes in cell composition. Using 10X single-cell sequencing technology, we performed single-cell transcriptome analysis of PBMCs from six young people aged 20–29 years and six elderly people aged 80–88 years. We sequenced 127,707 cells with an average of 10,642 cells per sample (Extended Data Table S5). We combined single-cell transcriptional profiles collected from the different samples for UMAP-based cell cluster analysis and annotated specific genetic markers for the resulting cell types. A total of 18 different cell clusters were identified (Figure 5A, Figure S3A, and Figure S3B). We then performed UMAP clustering on samples A and Y (Fig. 5B), calculated the relative percentages of the 18 major cell types in each person’s PBMC based on the single-cell RNA-seq (scRNA-seq) data (Fig. 5C), and quantified the percentage changes in cell types driven by aging (Fig. 5D). We found that the proportions of CD8+ T cells, T helper cells, B cells, and erythrocytes were significantly decreased in the older samples (p < 0.05), while the proportions of effector CD8+ memory T cells were significantly increased. This is basically consistent with the results of other studies^14^. The decline of B cells, T cells, and CD8+ T cells with age leads to the decline of T cell functions and B cell functions, and ultimately leads to the decline of human immune function. On the other hand, CD8+ memory T cells, which are associated with cytotoxicity and chronic inflammatory responses, increased.

**Figure 5.**
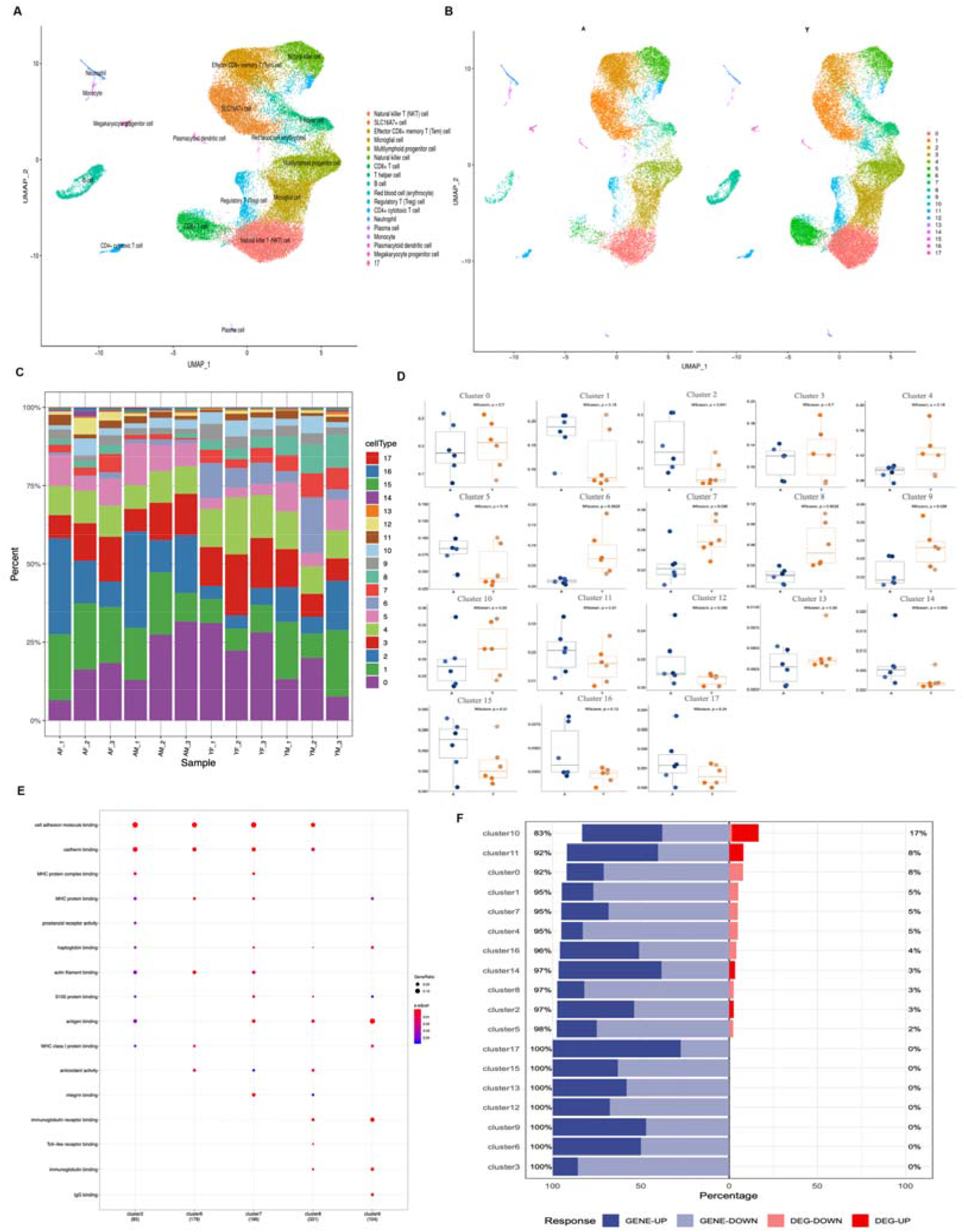
Characterization of immune cells in individuals across two conditions. **A** Identification of cell subpopulations from all analyzed samples. Samples from Y (n = 6) and A (n = 6), a total of 12 samples, showing 18 clusters with their own labels. Each point corresponds to a cell, colored according to the cell type. **B** UMAP projections of group A and group Y. Each point corresponds to a cell, colored according to the cell type. **C** The average proportion of each cell type from the 12 samples. **D** The proportion of each cell type in each sample was stained by the source donor. The x-axis corresponds to the grouping of samples (A, n = 6; Y, n = 6). The exact p-values on both sides that pass the Wilcoxon rank sum test are shown. Minimum whisker, 25th - 1.5 × inter-quartile range (IQR) or the lowest value within; minimum box, 25th percentile; center, median; maximum box, 75th percentile; maximum whisker, 75th percentile + 1.5 × IQR or greatest value within. **E** Differential gene analysis (p < 0.05, logFC > 0.1) was performed on group A and group Y of five cell subpopulations of clusters 2, 6, 7, 8, and 9. The top 20 enriched biological processes were obtained by GO analysis of the differential genes. The dot color indicates the statistical significance of enrichment (p), and the dot size indicates the percentage of genes annotated for each term. **F** Raised and lowered genes from the 18 clusters compared with the 787 genes, genetic variations from which to choose 18 clusters (|logFC| > 0.1, p < 0.05). Groups are shown in proportion.

The level of humoral immunity in the elderly was elevated. Through the detection of cytokines, we found that various inflammatory factors involved in humoral immunity were elevated in the elderly (Figure S3C), including TNF-α and IL6. On the other hand, GO enrichment analysis of the differential genes showed that CD8+ T cells, T helper cells, B cells and effector CD8+ memory T cells in four clusters were enriched in cell adhesion molecule binding (p < 0.05), and CD8+ memory T cells in four clusters were enriched in erythrocyte enrichment to antigen binding (p < 0.05) (Fig. 5E). Then we determined whether the top age-related genes in the batch RNA-seq were widely expressed in a single cell type or in multiple cell types. By comparing the 18 cell clusters with the 787 genes in the gene cluster (|logFC| > 0, q < 0.05), and the differences between the genes in the 18 clusters (|logFC| > 0.1, q < 0.05), we found that the 787 genes rarely differed from those in the 18 clusters (Fig. 5F, Extended data Table S6). This proves that the gene expression changes observed are caused by internal changes to the cells induced by age, rather than changes in the cell composition.

### Aging prediction model based on the transcriptome

We then built a model based on the above results that predicts the actual age based on the number of expressed genes in healthy individuals. We used a penalty regression model (elastic net) to regress the previously obtained z-score values of the 787 genes and the ages of the 113 samples. The elastic net regression model automatically selected 46 genes (penalty terms, α = 0.92, λ = 1). We called these 46 genes the “transcriptome clock” because their weighted average (formed by regression coefficients) is equivalent to a clock marking biological age. To predict the accuracy of the model, we used the Pearson correlation coefficient between the predicted age and the actual age, the MAE, and the regression coefficient R^2^ for linear regression of the predicted age and the actual age. From this we can see that the age prediction model performs very well (age correlation = 0.95, MAE = 5.203, R^2^ = 0.9) (Figure 6A, Extended data Table S7). The accuracy of this model is significantly better than Fleischer’s transcriptome data based on human dermal fibroblasts (R^2^ = 0.81, MAE = 7.7) (Fleischer *et al*. 2018). In addition, we also used these 787 sites to establish our own methylation age prediction model (Figure S4A, Extended data Table S8). A total of 34 genes were selected, and the model also performed very well (α = 1, λ = 0.94, age correlation = 0.99, MAE = 3.28, R^2^ = 0.97), better than Horvath’s methylation age model (age correlation = 0.96, MAD (Mean Absolute Deviation) = 3.6 years) (Horvath 2013).

**Figure 6.**
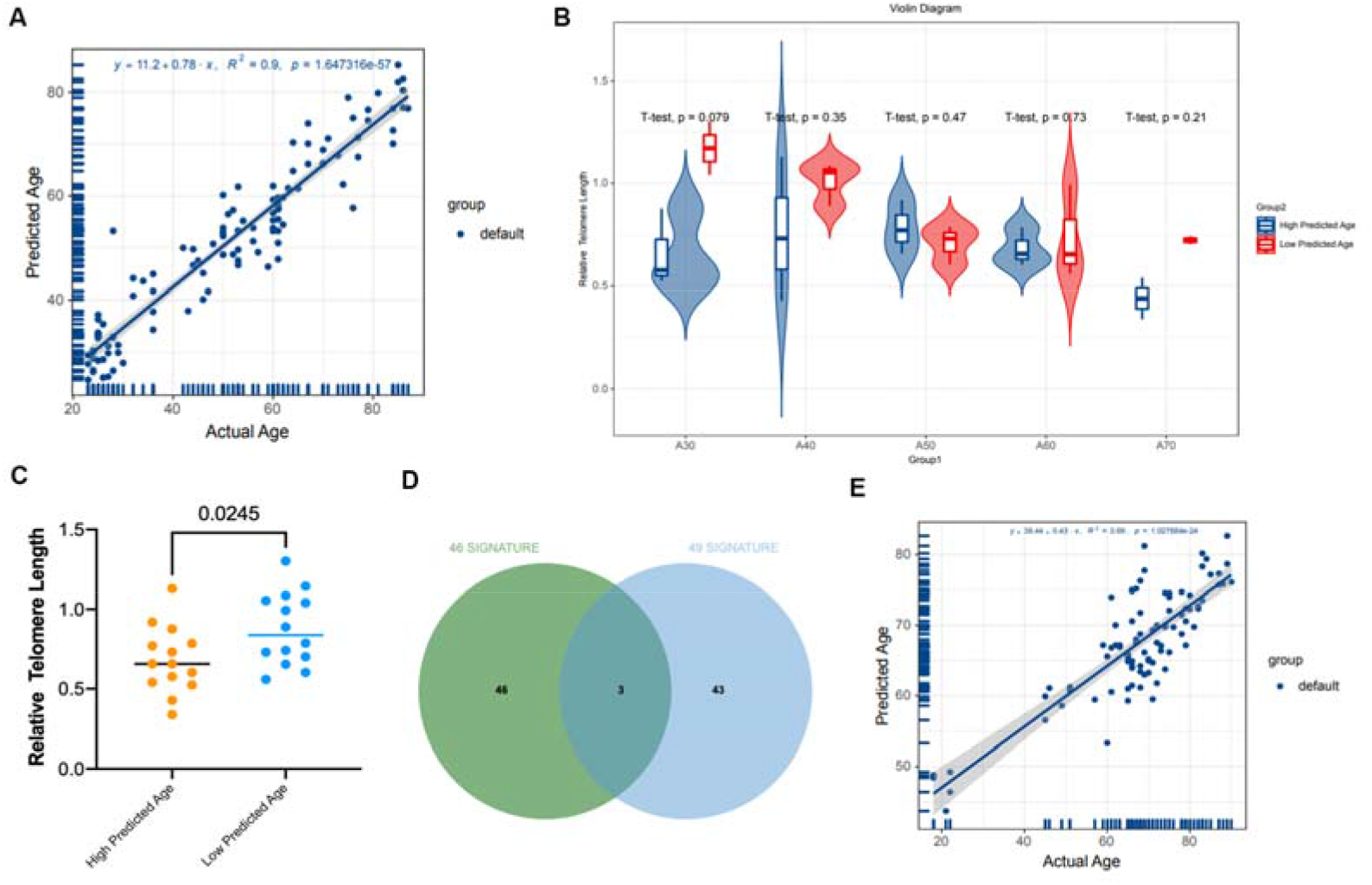
The aging clock models. **A** Cross-validation of 113 transcriptome samples based on 787 genes to predict age. Each point represents a transcriptome sample, showing the best-fit line between the predicted age and the true age, and the shading shows the 95% confidence interval of the line determined by resampling. The text at the bottom of each panel shows the performance indicators of goodness of fit such as the MAE and the R^2^ Pearson correlation coefficient of the best fit line. **B** Detection of the relative telomere length of individuals with high predictive age and individuals with low predictive age in each age group in the transcriptome aging clock (two-tailed t-test). **C** Comparison of relative telomere length between high and low predicted age groups (two-tailed t-test). **D** Comparison of the 46-gene signature based on our data with the 49-gene signature based on data from three independent cohorts. **E** Cross-validation of 91 transcriptome samples based on three independent cohorts to predict age. Each point represents a transcriptome sample, showing the best-fit line between the predicted age and the true age, and the shading shows the 95% confidence interval of the line determined by resampling. The text at the bottom of each panel shows the performance indicators of goodness of fit, such as the MAE and the R^2^ Pearson correlation coefficient of the best fit line.

To further verify the accuracy of the model, we tested the telomere length of the relevant samples. Telomere length is currently recognized as a marker of aging. In the five age groups of 30, 40, 50, 60, and 70 years old, we selected three people whose predicted age was greater than their actual age (rapid aging), and three whose predicted age was less than their actual age (slow aging), and performed DNA detection of the telomere lengths. The results were very surprising. Because of the problem of sample size, there was no significant difference between the two age groups (Figure 6B). It can be seen that the telomere length of the slow-aging group is relatively larger than that of the fast-aging group. If all the samples are combined, we can see that the telomere length of the population whose predicted age is less than their actual age is significantly higher than that of the population whose predicted age is greater than their actual age (p = 0.0245) (Figure 6C). In the methylation age model, because of the small number of samples, we only detected the telomere length of eight samples, showing similar trends, but not significant (p = 0.1232) (Figure S4B). These results show that our model can be accurate and can predict the accelerated aging of individuals.

To assess whether these findings are representative of the general population, we decided to validate our results using sequencing data from other cohorts. However, unfortunately, no public data that matched our data with both the same sequencing platform (DNBSEQ) and the same sample type (PBMCs) could be found in the public database (GEO). We ultimately compared our results with transcriptomic data of three independent cohorts from China, USA, and Europe, of which all the sequences were obtained from PBMCs. The total sample size of the three cohorts was 91 with participants ranging in age from 18 to 90 years old, although some patients had the chronic diseases of coronary heart disease and Alzheimer’s disease. Although there was a large overlap between data from these independent cohorts and our data, and 80% of the data could be covered by data from the other three independent cohorts, because of the sequencing platform (GSE153100: Illumina NovaSeq 6000; GSE153104: Illumina HiSeq 2500; GSE166780: HiSeq X 10), differences in sequencing depth and regional ethnicity resulted in many independent genes among the four datasets (Figure S4C). In the validation, we constructed a 46-gene signature with multiple genes missing in these three cohorts, which led to our inaccurate prediction results. So, we tried to re-run the 766 genes in these three cohorts that overlapped with the 787 genes we screened to observe the labeling differences under the same method by performing elastic network regression analysis, and we still chose the same conditions as before (penalty terms, α = 0.92, λ = 1) to obtain a 49-gene signature (Figure 6D). We compared the 49-gene signature with our previous 46-gene signature, and we found that only three genes were common; moreover, the prediction model for this data did not perform very well for age prediction (age correlation = 0.83, MAE = 7.39, R^2^ = 0.69). This is associated with the distribution of the data, with most of the samples existing in the 60– 90 age range and only five samples distributed in the 18–45 age range, which led to a decrease in the accuracy of the model. Also, there was a huge batch effect between the three datasets (Figure S4D), and although we used COMBAT to remove the batch effect, this also resulted in the data itself being altered, which is another of the reasons that it was difficult to use this data for model testing.

## Discussion

Our research spans the transcriptome of PBMCs, DNA methylation, a partial plasma proteome, and single-cell RNA-seq time-resolved information throughout the human adult life cycle. Previous studies have found that men and women age differently. In plasma proteomics, two-thirds of the proteins that change with age also change with sex (895 of 1,379 proteins) (Lehallier *et al*. 2019). In our research, we found that the differential genes between the sexes also differed with age.

By analyzing the aging transcriptome, we have determined the fluctuations during human life. These changes are the result of expression clusters moving in different patterns. We have provided two linearly correlated waves of the aging transcriptome (Figure 2C), with little overlap between the two key biological pathways. The correlation analysis between these two clusters and DNA methylation showed that the linear fluctuation of most genes with aging was regulated by DNA methylation in the promoter region of the related genes. However, in relation to the proteomics, we found that few differential proteins were regulated by the transcriptome, which may be related to the low detection efficiency of the protein sequencing. We only obtained 1,377 proteins, of which only 103 were differential proteins. We ended up with 787 genes that fluctuate with age. Single cell transcriptome sequencing showed that the gene expression changes were caused by internal cellular changes induced by age rather than changes to the cellular composition.

Finally, we built a regression model from the transcriptome data that allowed us to accurately predict the age of an individual. The mean absolute error in predicting age to five years is better than previous methods used to predict age based on genomic biomarkers. The application of this predictive approach in longitudinal studies raises the possibility of developing tools for monitoring and predicting aging and related diseases and provides tools for evaluating interventions to reverse aging.

It is also important to verify if the model can be confirmed using histological data generated using a different platform; however, thus far we have only found negative results. Therefore, we need to wait for data from the same platform to validate our results. In conclusion, in this study, 787 age-associated conserved genes were identified. Examination of the functions of these genes individually or in common may further reveal the underlying mechanisms of the aging process. Finally, age-based transcriptomic profiling of 46 gene expression profiles may capture the rate of aging and may help identify individuals who are aging faster than their actual age.

## Methods

### Ethics approval and consent to participate

This study was approved by Medical Ethics Committee of Tongji University Affiliated Shanghai East Hospital (2020-069). The participants provided their written informed consent to participate in this study.

### Research cohort

The 139 healthy individuals in this study were recruited to the study from Shanghai East Hospital (ages 23–88 years old; median, 53 years old). All the participants provided written consent. Participants were excluded if they had diabetes, advanced liver disease or renal failure, class III/IV congestive heart failure, myocardial infarction, unstable angina, stroke, uncontrolled hypertension or hyperglycemia, drug or alcohol abuse, had a history or presence of malignant tumors, had active autoimmune diseases, had any active infection (including positivity for hepatitis BsAg, hepatitis C antibody, or HIV antibody, or a positive PPD test). Three separate cohorts (GSE153100, GSE153104, and GSE166780) are available in GEO (Dhanwani *et al*. 2020; Wahl *et al*. 2021; Wang *et al*. 2021).

### Blood sample collection and processing

Between 7:00–9:00 in the morning, 6 ml of peripheral blood of the subject was drawn with a disposable vacuum negative pressure blood collection tube containing EDTA. The samples were centrifuged at 1000 × *g* for 10 min to separate plasma. After separating the plasma, the remaining blood cells were added to an equal volume of PBS and mixed gently upside down. We added 4 ml of lymphocyte separation solution to a 15 ml centrifuge tube and centrifuged it at 400 × *g* for 25 min. After centrifugation, stratification could be seen. The middle layer was slowly aspirated and transferred to a new 15 ml centrifuge tube, 3 ml PBS was added, and the sample was centrifuged at 600 × *g* for 10 min. Then the supernatant was discarded, 3 ml PBS was added, and it was centrifuged at 600 × *g* for 5 min. The supernatant was discarded and the cells were aliquoted (they were stored separately for DNA, RNA, and single cell sequencing). We added 800 μl Trizol to the RNA sample, whereas for single cell sequencing we used 90% FBS and 10% DMSO, and the samples were all stored at −80°C.

### DNA extraction

The sample was thawed, then centrifuged at ~11,200 × *g* for 1 min, and the supernatant was discarded. Using a Tiangen kit (Tiangen, Beijing, China), 200 μl of GA buffer was added and shaken, then 20 μl of Proteinase K solution was added and shaken gently to mix. We added 200 μl of GB buffer and mixed well, then placed at 70°C for 10 min, and then centrifuged briefly to remove water droplets on the inner wall. We then added 200 μl of absolute ethanol and shook to mix. The solution obtained in the previous step was added to a CB3 adsorption column placed in the collection tube and centrifuged at 11,200 × *g* for 30 s. The waste liquid was discarded, and the adsorption column was put back into the collection tube. About 500 μl of GD buffer was added to the adsorption column, it was centrifuged at 11,200 × *g* for 30 s, the waste liquid was discarded, and the adsorption column was returned to the collection tube. We added 600 μl of PW rinsing solution to the CB3 adsorption column, centrifuged again at 11,200 × *g* for 30 s, discarded the waste liquid, and put the adsorption column back into the collection tube. The previous step was repeated. The adsorption column was put back into the collection tube, centrifuged again at 11,200 × *g* for 2 min, and the waste liquid was discarded. The CB3 adsorption column was then left at room temperature for 5–10 min for the residual rinsing solution in the adsorption column to dry. The CB3 adsorption column was then put into a clean centrifuge tube, and 50–200 μl ddH2O was added to the middle of the adsorption membrane with a pipette. The column was stood at room temperature for 2–5 min, centrifuged at 11,200 × *g* for 2 min, and the solution was collected into the centrifuge tube. The RNA was then quantified with a NanoDrop (NanoDrop Technologies, Delaware, USA) and frozen at −80°C.

### 850K methylation sequencing

A SMA 400 UV-vis (Merinton, Shanghai, China) DNA/Proteins Analyzer was used to determine the purity and crude concentration of the samples. The concentration of the sample was measured using a Qubit 3.0 Fluorometer. The integrity of the samples was determined by agarose gel electrophoresis. Using a Zymo Research EZ DNA Methylation Kit, bisulfite transformation of genomic DNA was conducted to change the methylated cytosines and convert unmethylated cytosines into uracil. An Illumina Infinium Methylation EPIC Bead Chip Kit was used to detect the DNA transformed by bisulfite, and determined the methylation of CpG sites. The chip raw data was imported into the R package Champ (V2.12.4) to calculate the raw beta values, and the PBC method in the Champ package was used for standardization. Referring to the genomic location, different methylation sites were found at the same location in the genomes of multiple samples. The two groups of samples were first standardized, then the linear regression method was used to detect the different methylation sites, and the sites with a p-value < 0.01 were screened as different methylation sites.

### RNA extraction

To prepare the samples, the Trizol lysate was replenished to 1 ml in the sample tube, pipetted repeatedly, and let stand for 5 min at room temperature. It was then centrifuged at 12,000 × *g* for 5 min at 4°C. The supernatant was transferred to a sterilized 1.5-ml centrifuge tube, 0.3 ml chloroform was added, it was shaken for 15 s and then let stand at room temperature for 5 min. The sample was then centrifuged at 12,000 × *g* for 10 min at 4°C. After centrifugation, the liquid phase had separated and the liquid was divided into three layers: the bottom layer was the organic phase, and the middle and upper layers were both phases of water containing RNA. The upper aqueous phase was slowly transferred to a new 1.5 ml tube and an equal volume of isopropanol was added, the sample was vigorously mixed upside down and then let stand at −20°C for 2 h. The sample was centrifuged at 11,200 × *g* for 20 min at 4°C, and the precipitated RNA formed the pellet. The supernatant was discarded, the RNA pellet was resuspended and washed in 1 ml of 75% ethanol by flicking the tube wall, and centrifuged at 11,200 × *g* at 4□ for 3 min. The previous step was repeated and then the ethanol was carefully discarded and the RNA pellet was dried. The RNA precipitate was then dissolved in 25–100 μl DEPC water and the concentration was checked. The integrity of the RNA sample was checked with 1% agarose electrophoresis and 1 μl was used to measure the OD_260_/OD_280_ ratio on a spectrophotometer to detect the purity of the sample. The rest of the RNA was stored at −80□.

### cDNA synthesis, library construction and RNA sequencing

Temperature-suitable denaturation of the RNA sample was performed to open its secondary structure, and then oligo (dT) magnetic beads were used to enrich the mRNA. Reagents were added that interrupt the RNA sample, so the mRNA can be fragmented. The already configured first-strand synthesis reaction was added to the fragmented mRNA, then the first-strand cDNA was synthesized on a PCR machine. The second-strand synthesis reaction system was prepared again, and after a certain temperature reaction, two-strand cDNA was synthesized.

The reaction system was configured to repair the ends of the double-stranded cDNA, add A bases to the 3’ end, prepare a linker connection reaction system, and react at a suitable temperature for a certain period of time so that the linker is connected to the cDNA. The PCR reaction system was configured, the reaction program was set, and the product was amplified. After denaturing the PCR product into a single-stranded product, a DNA circularization reaction system was prepared, the sample was fully shaken and mixed, and reacted at a suitable temperature for an appropriate time to obtain a single-stranded circular product. The uncircularized linear DNA molecules were then digested to form the final library construction. Single-stranded circular DNA molecules can be replicated through rolling circles to form a DNA nanosphere (DNB) containing more than 200 copies. Afterwards, the obtained DNBs were added to the mesh holes on the chip using high-density DNA nanochip technology. Then, using combined probe anchor polymerization technology (cPAS) for sequencing, we obtained sequencing read lengths of 50 bp/100 bp/150 bp. In this study, the DNBSEQ platform was used to sequence the samples. Before data analysis, reads with low quality and high positional base N content were first removed to ensure the reliability of the results. Then Bowtie2 was used to align the clean reads to the reference sequence (Kim *et al*. 2015), and the RSEM tool was used to calculate the expression level of the gene and transcription sequence (Li and Dewey 2011). Finally, we used the RStudio script and the following Bioconductor packages for data visualization and analysis: Rtsne, Deseq2, topGO, destination, and org.m.eg.db. For differential gene screening, the screening conditions were artificially set as follows: fold change (FC) ≥ 1 and p-value < 0.05.

### Protein extraction and data-independent acquisition (DIA) quantification

#### Protein extraction

SDS-free L3 was added to 100 μl of blood sample to make the volume up to 1 ml; then a reductive alkylation reaction was conducted. The specific steps were as follows: DTT (dithiothreitol) was added to make the final concentration 10 mM, the sample was placed in a 37□-water bath for the reaction to occur for 30 min, and then placed at room temperature once the reaction was over. IAM (iodoacetamide) was added immediately after the temperature was dropped to room temperature to a final concentration of 55 mM, and the reaction was conducted in a dark room at room temperature for 30 min. The processed protein solution was then enriched by solid phase extraction with a C18 column. A new C18 column was taken and the sample was passed through the column with 1 ml of methanol at a flow rate of 3 drops/s. It was then passed through the column with 1 ml of 0.1% FA at a flow rate of 3 drops/s; if the sample volume was less than 1 ml, it was diluted to 1 ml with SDS-free L3 and passed through the column, while if the sample volume was greater than 1 ml, it could be passed through the column in multiple parts. The sample was loaded twice at a flow rate of 1 drop/s, using 1 ml 0.1% FA to pass the column at a flow rate of 3 drops/s, this step was repeated three times in total. Then 800L 75% acetonitrile (ACN) was used to slowly elute with a flow rate of 0.5 drop/s, and the eluent was freeze-dried before proceeding to the next step.

#### Peptide separation

Using the Shimadzu LC-20AD liquid phase system, 20 μg of sample was taken after mixing. The separation column used was a 5-μm 20-cm × 180-μm Gemini C18 column, and the sample was separated in liquid phase using the separation column. Then mobile phase A (5% ACN, pH 9.8) was used to redissolve the dried peptide sample and load the sample, and then gradient elution was performed at a flow rate of 4 μl/min in the following steps: 2% mobile phase B (95% ACN, pH 9. 8) elution for 7 min, 2% to 7% mobile phase B for 3 min, 7% to 25% mobile phase B for 27 min, 25% to 60% mobile phase B for 2 min, 60% to 80% mobile phase B for 1 min, mobile phase B for 3 min, 80% to 2% mobile phase B for 2 min, and then equilibrated in 2% mobile phase B for 5 min. The elution peak was monitored at a wavelength of 214 nm and a component was collected every 3.15 min, combined with the chromatographic elution peak figure to obtain ten components, and then freeze-dried.

#### High pH RP separation

The same amount of peptides were taken from each sample and mixed into a tube. The peptide content was 20 μg, the mixed 20 μg of peptides were diluted and loaded with 2 ml mobile phase A (containing 5% ACN, pH 9.8). Then, the Shimadzu LC-20AB liquid phase system was used, and the separation column used for liquid phase separation of the sample was a 5-μm 4.6- × 250-mm Gemini C18 column. The sample was then eluted with a flow rate gradient of 1 ml/min: it was first eluted with 5% mobile phase B (95% ACN, pH 9.8) for 10 min, and then eluted with 5% to 35% mobile phase B for 40 min, eluted with 35% to 95% mobile phase B for 1 min, and finally mobile phase B was used for continuous elution for 3 min, and then the sample was equilibrated with 5% mobile phase B for 10 min at room temperature. Then, wavelength detection was conducted at 214 nm. According to the elution peak, one component was collected every minute, and then the samples were combined according to the elution peak diagram of the chromatogram. Finally, components were obtained, and the ten components were drained and frozen.

#### Data-dependent acquisition (DDA) library construction and DIA quantitative detection (Nano-LC-MS/MS)

First the peptide sample that was freeze-dried in the previous step was dissolved with mobile phase A (containing 2% ACN and 0.1% FA) and centrifuged at 20,000 × *g* for 10 min. The supernatant was aspirated and the sample was loaded. The UltiMate3000 UHPLC (Thermo Fisher, Sunnyvale, CA, USA) was used for preliminary separation of the samples. The sample was added to the trap column for enrichment and desalination, and then the trap column was connected in series to a self-assembled C18 column (the inner diameter of the adsorption column was 150 μm, the particle size of the column material was 1.8 μm, and the column length was about 35 cm), and then the column was gradient separated at a flow rate of 500 nl/min. In the first step, 5% mobile phase B (98% ACN and 0.1% FA) was separated for 5 min, then mobile phase B was linearly increased from 5% to 25% for 90 min. Then, mobile phase B was increased from 25% to 35% for 10 min, then mobile phase B was increased from 35% to 80% for 5 min. The sample was maintained at 80% mobile phase B for 3 min, and then mobile phase B decreased from 80% to 5% for 30 s. Then mobile phase B was maintained at 5% for 6.5 min. After liquid phase separation the peptides need to pass through the nano ESI source first, and then enter the Fusion Lumos tandem mass spectrometer after ionization, and then the second step of the DDA detection is performed. The main settings were as follows: the voltage of the ion source was set to 2 kV; in the primary mass spectrum, the scanning range of the primary mass spectrum was 350–1500 m/z; the resolution of the mass spectrometer was set to 120,000; and the maximum ion injection time (MIT) was 50 ms. In the secondary mass spectrum, the secondary mass spectrum was set to HCD fragmentation mode; the fragmentation energy was set to 30; the resolution was set to 30,000; the ion implantation time was a maximum of 100 ms; and the dynamic rejection time was 30 s.

The peptides after liquid phase separation were ionized by the nano ESI source, and then detected in the DIA mode by a tandem mass spectrometer. A Fusion Lumos (Thermo Fisher Scientific, San Jose, CA) instrument was used to collect mass spectral data from the samples in DIA mode. The peptides and proteins were quantified by MSSTATS software. After that, the MSSTATS software package was used to complete the steps of error correction and normalization in the system for each sample. Then according to the set comparison group and linear mixed effect model, the differences of different proteins were evaluated. Two conditions, a fold change > 1.5 and a p-value < 0.05, were used as screening criteria for significantly different proteins.

### Single cell sequencing

The frozen single cell samples were placed on a PCR apparatus for cell lysis. The reverse transcription system was prepared, and the first cDNA was synthesized by SMART amplification. Kapa HiFi Hot Start DNA Polymerase was used to pre-amplify the cDNA. Agencourt AMPure XP magnetic beads purified the double-stranded cDNA products and an Agilent 2100 BioAnalyzer was used to test the quality of the amplified products. The qualified amplification products were constructed by the transposable enzyme method, and an Agilent 2100 BioAnalyzer was used to detect the fragment size and concentration of the library. After denaturing the library into a single chain, the cyclization reaction system was prepared, and the single chain ring product was obtained by fully mixing the system for a certain temperature reaction time. Enzyme digestion of uncycled linear DNA molecules was conducted, followed by quantitative cyclization of libraries using a Qubit Fluorometer. Single-stranded circular DNA molecules were replicated by rolling rings to form a DNA nanosphere (DNB) containing more than 200 copies. The resulting DNBs were added into the mesh holes on the chip using high-density DNA nanochip technology. Sequencing was performed using combinatorial probe-anchor-anchor synthesis (CPAS).

The filtered gene expression matrix package (v.3.0.0) was analyzed by R software (v.3.5.3) and Seurat 43. Genes expressed at a data ratio of > 0.1% and cells with > 200 genes detected were selected for further analysis. Low quality cells were removed if the following criteria were met: (1) < 800 unique molecular identifiers (UMIs); (2) < 500 genes; or (3) > 10% of the UMIs were from the mitochondrial genome. After the low-quality cells were removed, the gene expression matrix was normalized by the Normalize Data function and 2,000 features with high intercellular variability were calculated using the FindVariableFeatures function. To reduce the dimensionality of the data set, the runPCA function used the default parameters to scale the linear transformation data generated by the scaleData function. Next, elbowPlot (as suggested by the Seurat developers), the DimHeatMap, and JackstrawPlot functions were used to identify the true dimensions of each dataset. Finally, we used the findNeighbors and findClusters functions to cluster the cells, and the runuMap function, which is set by default, to perform non-linear dimension reduction. After the non-linear dimensional reduction of all cells by UMAP and the projection into a two-dimensional space, the cells were clustered together according to common characteristics. The findAllMarkers function in Seurat was used to find markers for each identified cluster. Clusters were then classified and annotated according to the expression of typical markers for a particular cell type. The differential gene expression test was performed using the findMarkers function in Seurat, with the default parameter “test.Use = Wilcox”, and the Benjamini-Hochberg method was used to estimate the rate of false discovery (FDR). The DEGs were filtered with a minimum log_2_(multiple change) of 0.5 and a maximum FDR value of 0.01.

### Telomere length detection

#### Telomere gene PCR system and reaction process

The reaction process included pre-denaturation at 95°C for 10 min, denaturation at 95°C for 15 s, followed by extension at 54°C for 2 min, and 23 cycles in the last two steps.

#### 36B4 internal reference gene PCR system and reaction process

The reaction process was pre-denaturation at 95°C for 10 min, denaturation at 95°C for 15 s, followed by extension at 58°C for 1 min; the last two steps were 33 cycles.

#### Primer sequences

Tel-F: 5’-GGTTTTTGAGGGTGAGGGTGAGGGTGAGGGTGAGGGT-3’.
Tel-R: 5’-TCCCGACTATCCCTATCCCTATCCCTATCCCTATC-3’.
36B4-F: 5’-CAGCAAGTGGGAAGGTGTAATCC-3’.
36B4-R: 5’-CCCATTCTATCATCAACGGGTACAA-3’.

### Statistical analysis

Unless noted otherwise, statistical analysis was performed using R, a language and environment for statistical computing and graphics.

### Age prediction model

The penalty-regression model (implemented in R-packet glmnet) was used to regress the time series by age of 787 genes (transcriptome penalty terms, α = 0.92, λ = 1; methylation penalty terms, α = 1, λ = 0.94).

### Data access

Raw sequencing data of RNA-seq, proteomics and Single-cell RNA seq were deposited into the CNGB Sequence Archive (CNSA) of the China National GeneBank Database (CNGBdb) with accession numbers CNP0002176. The source data of methylation are provided in the Extended Data (Extended Data Table S8).

## Acknowledgements

We thank all participants. This work was supported by grants from National Key Research and Development Program of China (2020YFC2002800), the Fundamental Research Funds for the Central University (22120210584), and Major Program of Development Fund for Shanghai Zhangjiang National Innovation Demonstration Zone<Stem Cell Strategic Biobank and Stem Cell Clinical Technology Transformation Platform> (ZJ2018-ZD-004).

## Authors’ contributions

C. H. performed experiments. C. H., Y. Z and H.L. analyzed data. H. J., Z.L. and H.L. supervised the project. C. H. and H.L. designed the project and wrote the paper.

---

**Figure S1.**
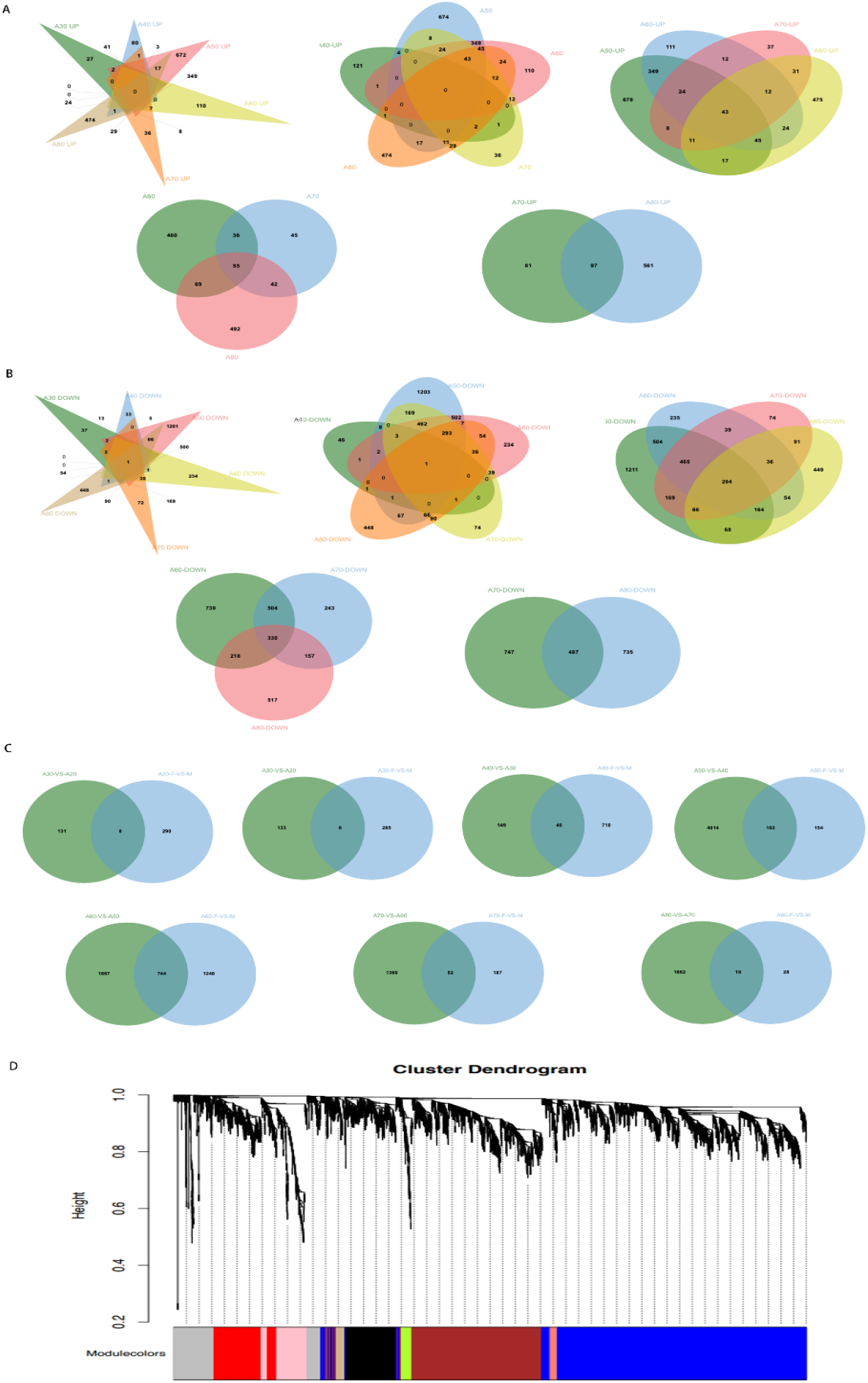
**A** Venn diagram for comparison of upregulated genes in the seven age groups. **B** Venn diagram of downregulated differential genes in the seven age groups. **C** Venn diagrams of age and sex differential genes. **D** Age-related modules in WGCNA, including 40 non-gray modules.

**Figure S2.**
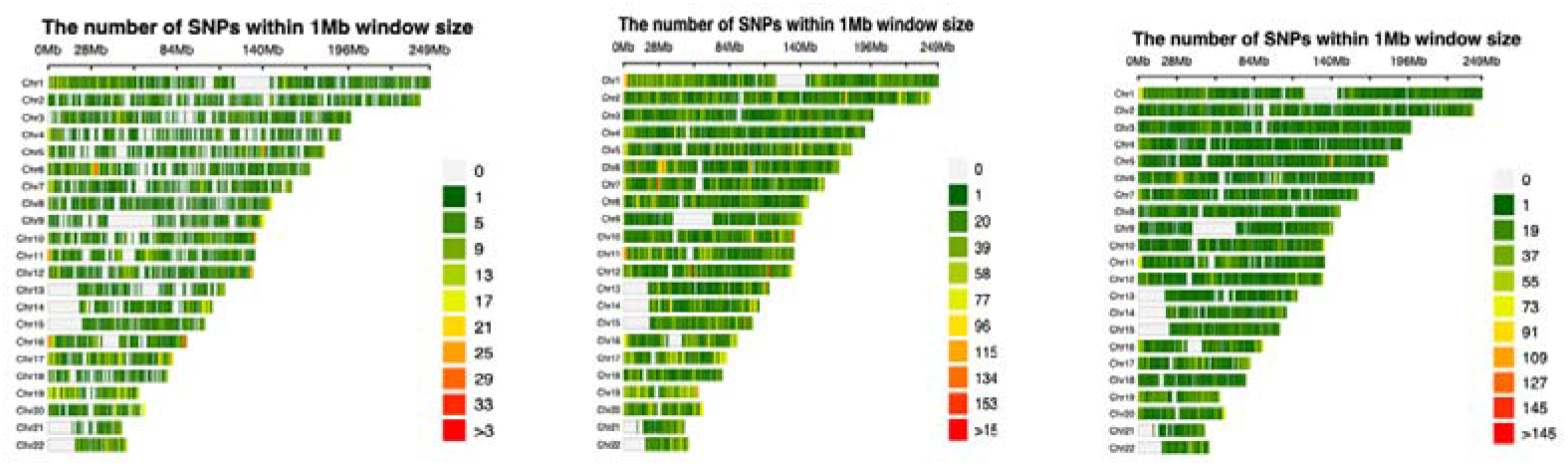
Distribution of differential methylation sites on chromosomes (groups M and Y, YO and Y, and A and Y).

**Figure S3.**
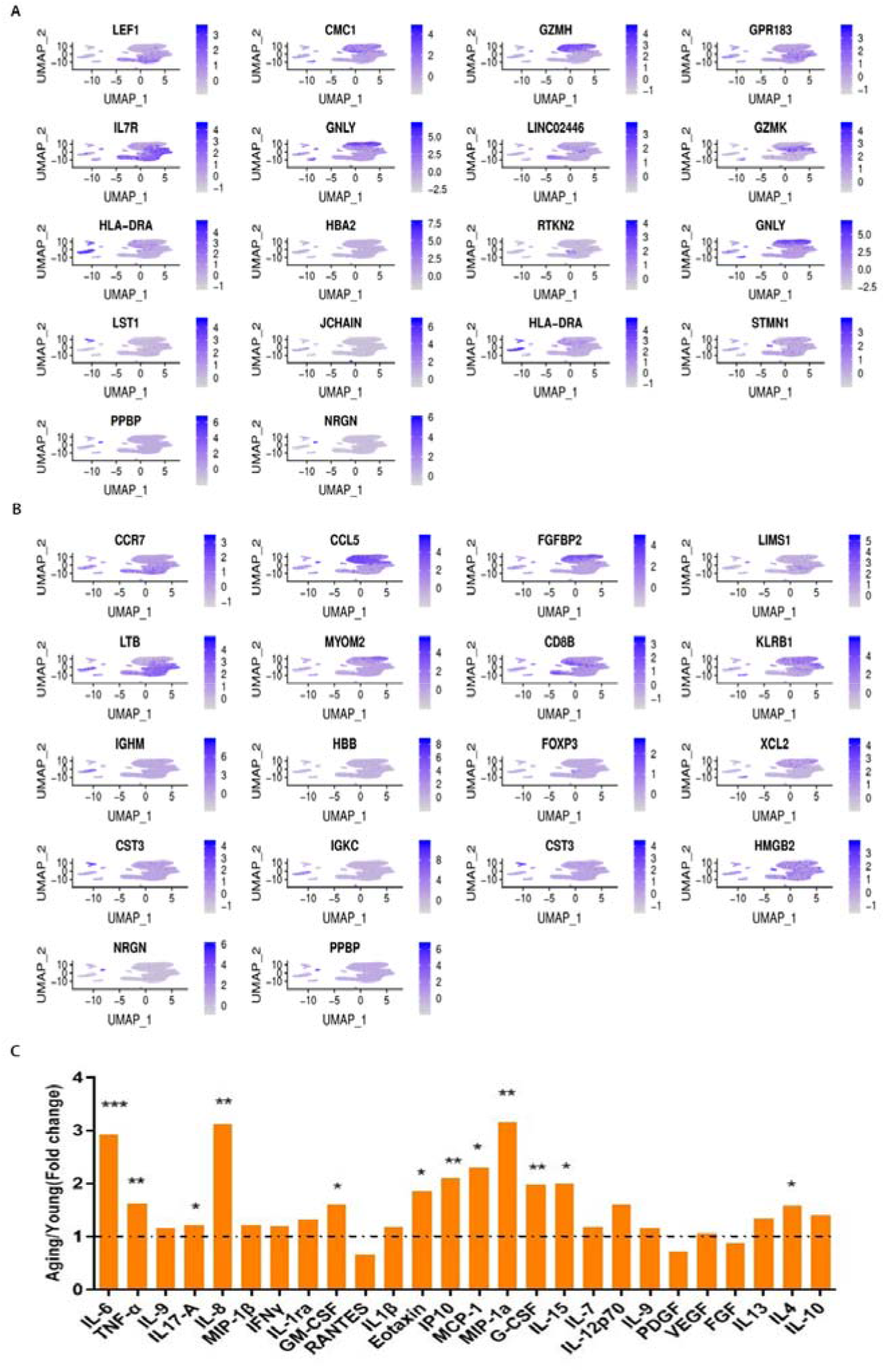
**A** Lattice heatmap of 18 cluster specific TOP1 marker genes. **B** Lattice heatmap of 18 cluster specific TOP2 marker genes. **C** Young age group compared with old age group for the levels of 26 kinds of cytokines (t-test; *, p < 0.05; **, p < 0.01; ***, p < 0.001).

**Figure S4.**
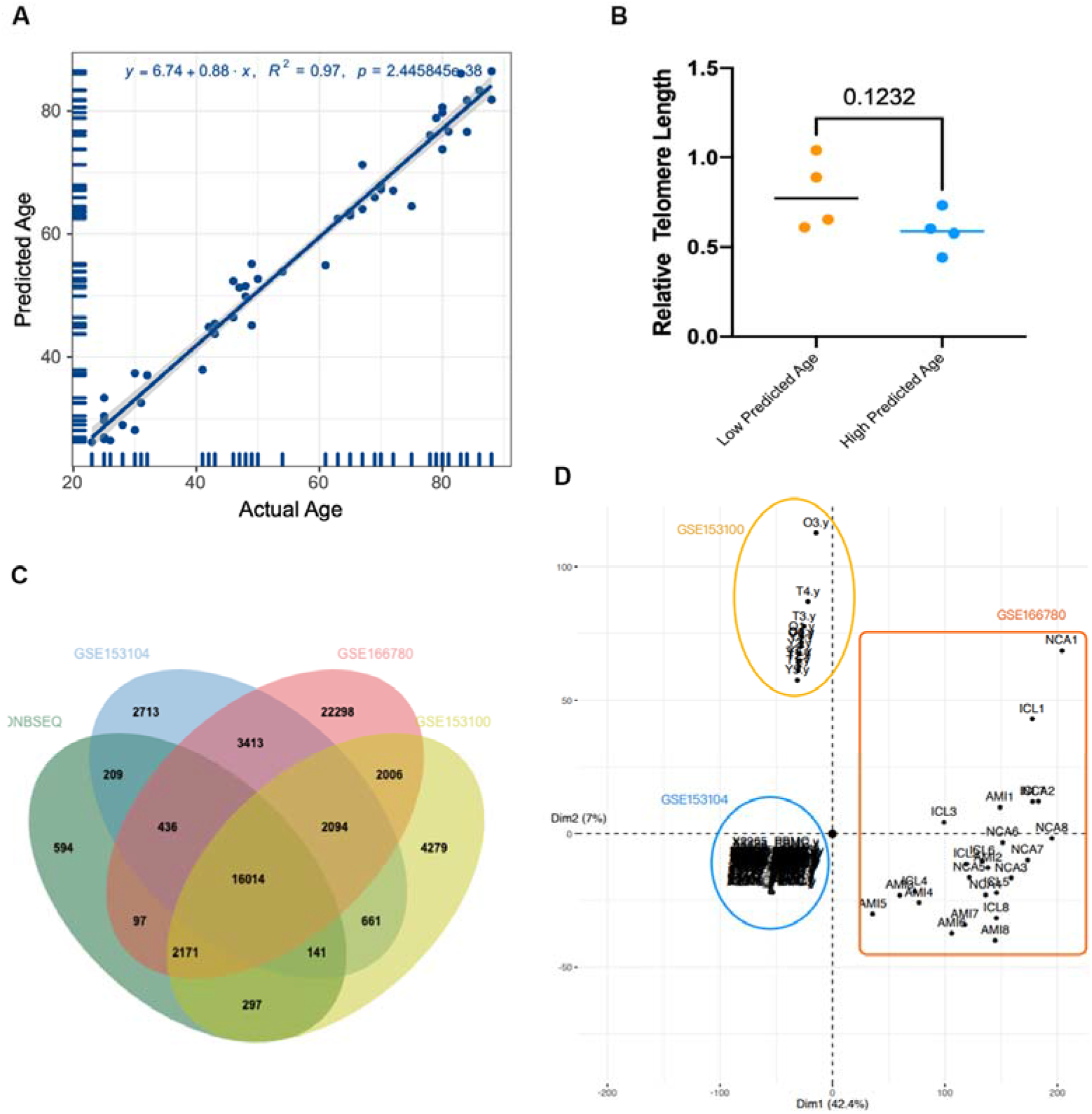
A Cross-validation of 49 DNA methylation samples based on 787 genes to predict age. Each point represents a transcriptome sample, showing the best-fit line between the predicted age and the true age, and the shading shows the 95% confidence interval of the line determined by resampling. The text at the bottom of each panel shows the MAE, the R^2^ of the best fit line, the Pearson correlation coefficient, and other performance indicators of goodness of fit. **B** Detection of the relative telomere length of some individuals with high predictive age and individuals with low predictive age in the methylation aging clock (two-tailed t-test). **C** Comparison of three independent cohorts and our sequencing data. **D** PCA analysis of the three cohorts.

